# The proBAM and proBed standard formats: enabling a seamless integration of genomics and proteomics data

**DOI:** 10.1101/152579

**Authors:** Gerben Menschaert, Xiaojing Wang, Andrew R. Jones, Fawaz Ghali, David Fenyö, Volodimir Olexiouk, Bing Zhang, Eric W. Deutsch, Tobias Ternent, Juan Antonio Vizcaíno

## Abstract

On behalf of The Human Proteome Organization (HUPO) Proteomics Standards Initiative (PSI), we are here introducing two novel standard data formats, proBAM and proBed, that have been developed to address the current challenges of integrating mass spectrometry based proteomics data with genomics and transcriptomics information in proteogenomics studies. proBAM and proBed are adaptations from the well-defined, widely used file formats SAM/BAM and BED respectively, and both have been extended to meet specific requirements entailed by proteomics data. Therefore, existing popular genomics tools such as SAMtools and Bedtools, and several very popular genome browsers, can be used to manipulate and visualize these formats already out-of-the-box. We also highlight that a number of specific additional software tools, properly supporting the proteomics information available in these formats, are now available providing functionalities such as file generation, file conversion, and data analysis. All the related documentation to the formats, including the detailed file format specifications, and example files are accessible at http://www.psidev.info/probam and http://www.psidev.info/probed.

## Introduction

Mass spectrometry (MS) based proteomics approaches have advanced enormously over the last decade, and are becoming increasingly prominent as an essential tool for post-genomic research. Proteomics approaches enable the identification, quantification and characterization of proteins, peptides, and post-translational protein modifications (PTMs) such as phosphorylation, providing information about protein expression and functional states [1]. Despite the instrumental role of the underlying genome in proteomics data analysis, it is only relatively recently when the field of proteogenomics started to gain prominence [2-4].

In proteogenomics, proteomics data is combined with genomics and/or transcriptomics information, typically by using sequence databases generated from DNA sequencing efforts, RNA-Seq experiments [5], Ribo-Seq approaches [6, 7], and long-non-coding RNAs [8], among others, in the MS-based identification process. Peptide sequences are mapped back to gene models *via* their genomic coordinates, demonstrating evidence of new translational events (e.g. novel splice junctions). Proteogenomics studies can be used to improve genome annotation and are increasingly utilized to understand the information flow from genotype to phenotype in complex diseases such as cancer [9-11] and to support personalized medicine studies [12].

Since 2002, the Proteomics Standards Initiative (PSI, http://www.psidev.info) of the Human Proteome Organization (HUPO) [13] has taken the role of developing open community standard file formats for different aspects of MS based proteomics analysis and data types. At present, well-established data standards are available for instance, for representing raw MS data (the mzML data format [14]), peptide and protein identifications (mzIdentML [15] and mzTab [16]) and quantitative information (mzQuantML [17] and mzTab).

The existence of compatible and interoperable data formats is a way to facilitate and advance “multi-omics” studies, and a clear need in proteogenomics, due to the growing importance of the field [9, 10, 18, 19]. However, no standard file format had been established so far for proteogenomics data exchange. To address this problem, we here present two novel standard data formats called proBAM and proBed. As suggested by their names, these two formats are adapted from their genomics counterparts BAM/SAM [20, 21] and BED (Browser Extensible Data) [22], where proBAM stands for proteomics BAM file (compressed binary version of the Sequence Alignment/Map (SAM) format) and proBed stands for proteomics BED file. A key feature of these formats is that they can seamlessly accommodate both regular genomic mapping information and specifics related to proteomics data, i.e. peptide-to-spectrum matches (PSM) or peptide sequence information. Existing popular genomics tools as SAMtools [20, 21] and Bedtools [23, 24], or the most widely used genome browsers such as Ensembl [25], the University of California Santa Cruz (UCSC) Genome Browser [26], JBrowse [27] and the Integrative Genomics Viewer (IGV) [28], can be used to manipulate and visualize proteomics data in these formats already. We believe that both proBAM and proBed are essential to merge the growing amount of proteomics information with the available genomics/transcriptomics data.

## Experimental Procedures

The development of these data formats has taken place since 2014 and it has been an open process *via* conference calls and discussions at the PSI annual meetings. Both format specifications have been submitted to the PSI document process [29] for review. The overall goal of this process, analogous to an iterative scientific manuscript review, is that all formalized standards are thoroughly assessed. This process is handled by the PSI Editor and external reviewers who can provide feedback on the format specifications. Additionally, there is a phase for public comments, ensuring the involvement of heterogeneous points of view from the community.

Both formats use Controlled Vocabulary (CV) terms and definitions as part of the PSI-MS CV [30], also used in other PSI data formats. All the related documentation, including the detailed file format specifications and example files, are available at http://www.psidev.info/probam and http://www.psidev.info/probed.

## Overview of the proBAM and proBed formats

The proteogenomics formats proBAM and proBed are designed to store a genome-centric representation of proteomics data (Figure 1). As mentioned above, both formats are highly compatible with their originating genomics counterparts, thus benefiting already from a plethora of existing tools developed by the genomics community.

**Figure 1.**
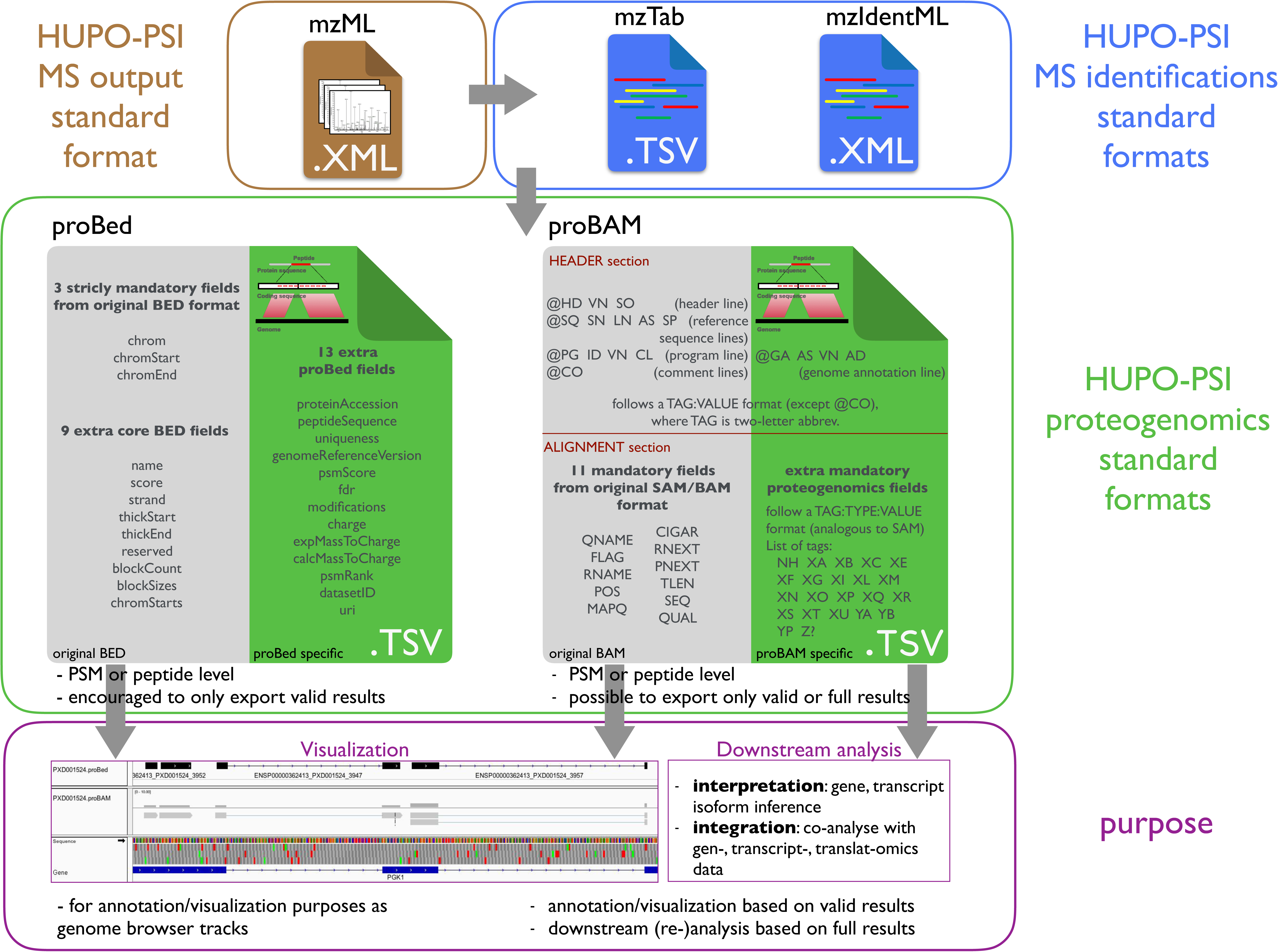
Overview of the proBAM and proBed proteogenomics standard formats. Both proBAM and proBed can be created from well-established proteomics standard formats containing peptide and protein identification information (mzTab and mzIdentML, blue box), which are derived from their corresponding MS-data spectrum files (mzML, brown box). The proBAM and proBed formats (green box) contain similar PSM related and genomic mapping information, yet proBAM contains more details, including enzymatic (protease) information, key in proteomics experiments (enzyme type, mis-cleavages, enzymatic termini, etc) and mapping details (CIGAR, flag, etc). Additionally, proBAM is able to hold a full MS-based proteomics identification result set, enabling further downstream analysis in addition to genome-centric visualization, as it is also the purpose for proBed (purple box).

### proBAM overview

The BAM format was originally designed to hold alignments of short DNA or RNA reads to a reference genome [20, 21]. A BAM file typically consists of a header section storing metadata and an alignment section storing mapping data (Figure 1, Figure 2 and Supplemental Table 1). The metadata can include information about the sample identity, technical parameters in data generation (such as library, platform, etc) and data processing (such as mapping tool used, duplicate marking, etc). Essential information includes where reads are aligned, how good the alignment is and the quality of the reads. Specific fields or tags are designed to represent or encode such information. The proBAM format inherits all these features. In this case, sequencing reads are replaced by PSMs (see proBAM specification document for full details, https://goo.gl/EW1cqB).

**Figure 2.**
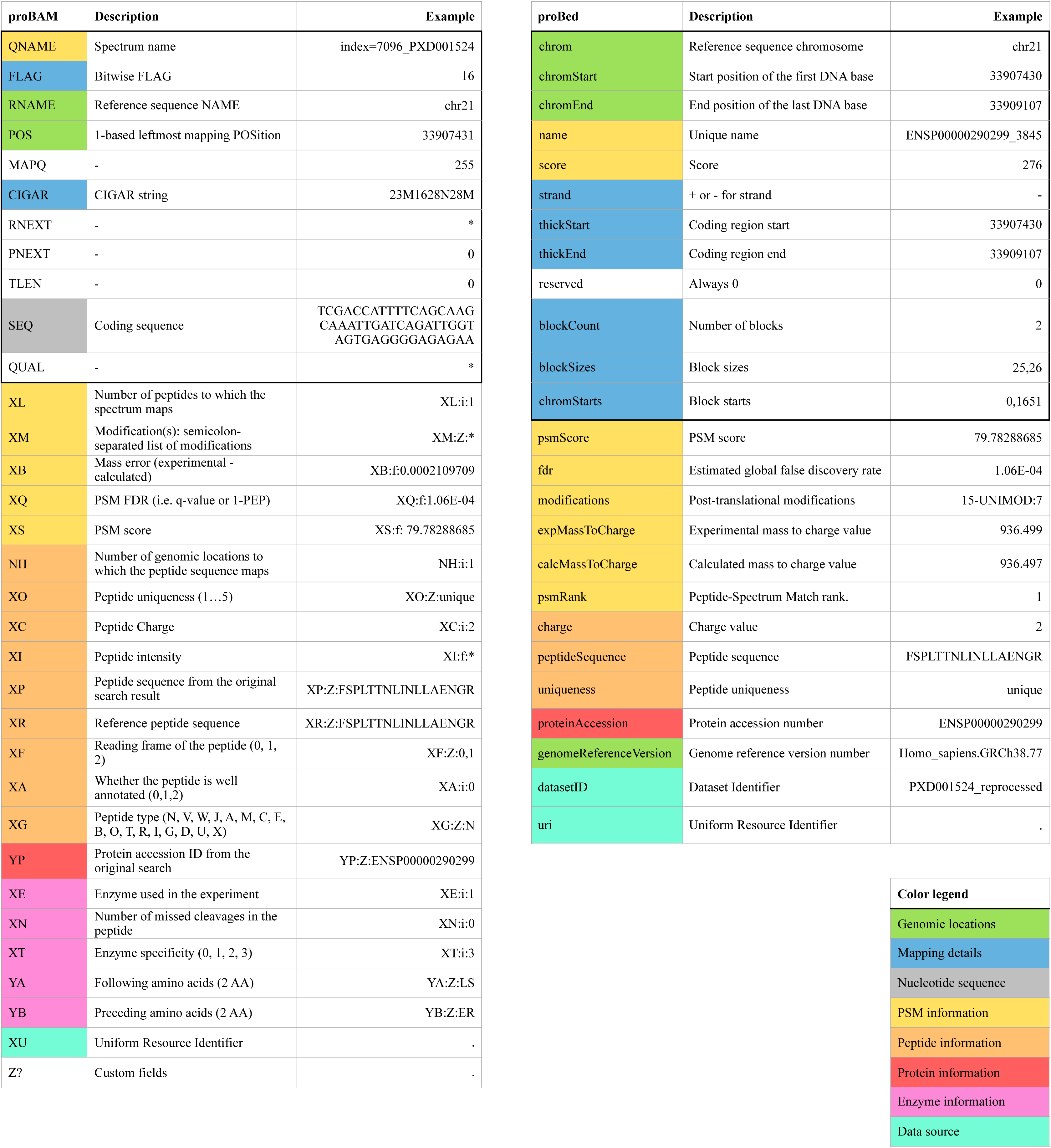
Fields of proBAM and proBed format. A proBed file holds 12 original BED columns (highlighted by a bold box) and 13 additional proBed columns. The proBAM alignment record contains 11 original BAM columns (highlighted by a bold box) and 21 proBAM-specific columns, using the TAG:TYPE:VALUE format. Each row in the table represents a column in proBAM and proBed. The rows are colored to reflect the categories of information provided in the two formats (see color legend at the bottom, the header section of proBAM format is not included here). The rows without any background color in the proBAM table represent original BAM columns that are not used in proBAM but that are retained for compatibility. The last row in grey indicates the customized columns that could be potentially used.

It should be noted that, since the tags used in BAM usually have recognized meanings, we did not attempt to repurpose any of them but rather created new ones to accommodate specific proteomics data types such as PSM scores, charge states, and protein PTMs (Figure 2 and proBAM specification document section 4.4.1 for full description on PSM specific tags). We also envisioned that additional fields and tags may be necessary to hold additional aspects of proteomics data. We thus designed a “Z?” tag as an extension anchor. Analogously to proBed, the format can also accommodate peptides (as groups of PSMs with the same peptide sequence). At the moment of writing, the proBAM format is under review as part of the PSI document process (http://www.psidev.info/probam). *As a note to editors and reviewers, we are aiming to conclude the PSI review at the same time as the manuscript describing proBAM/proBed is deemed suitable for publication, at which point we can announce a finalized standard.*

### proBed overview

The original BED format (https://genome.ucsc.edu/FAQ/FAQformat.html#format1), developed by the UCSC, provides a flexible way to define data lines that can be displayed as annotation tracks. proBed is an extension to the original BED file format [26]. In BED, data lines are formatted in plain text with white-space separated fields. Each data line represents one item mapped to the genome. The first three fields (corresponding to genomic coordinates) are mandatory, and an additional 9 fields are standardized and commonly interpreted by genome browsers and other tools, totalling 12 BED fields, re-used here. The proBed format includes a further 13 fields to describe information primarily on peptide-spectrum matches (PSMs) (Figure 1, Figure 2 and Supplemental Table 1). The format can also accommodate peptides (as groups of PSMs with the same peptide sequence), but in that case, some assumptions need to be taken in some of the fields (see proBed specification document Section 6.8 for details, https://goo.gl/FM2w66). At the moment of writing, the proBed format has completed the PSI internal review process, so the first version of the standard has been formalized (version 1.0, http://www.psidev.info/probed).

### Distinct features of proBAM and proBed and their use cases

The proBAM and proBed formats differ in similar ways as their genomic counterparts do, although representing analogous information. In fact, proBAM and proBed are complementary and have different use cases. Figure 3 shows two examples of proBAM and proBed visualization tracks of the same datasets. An IGV and Ensembl visualization are presented including multiple splice-junction peptides (Figure 3.A) and a novel translation initiation event in the HDGF gene locus (Figure 3.B), respectively.

Similar to the designed purposes of SAM/BAM, the basic concepts behind the proBAM format are: i) to provide genome coordinates as well as detailed mapping information, including CIGAR, flag, nucleotide sequences, etc; ii) to hold richer proteomics related information; and iii) to serve as a well-defined interface between PSM identification and downstream analyses. Therefore, the proBAM format contains much more information about the peptide-gene mapping statuses as well as PSM related information, when compared to proBed. Peptide and nucleotide sequences are inherently embedded in proBAM, which can be useful for achieving improved visualization by tools such as IGV. This feature enables intuitive display of the coverage of a region of interest, peptides at splice junctions, single nucleotide/amino acid variation, and alternative spliced isoforms (Figure 3), among others. Therefore, proBAM can hold the full MS proteomics result set, whereupon further downstream analysis can be performed: gene-level inference [31], basic spectral count based quantitative analysis, reanalysis based on different scoring systems and/or FDR (False Discovery Rate) thresholds.

**Figure 3.**
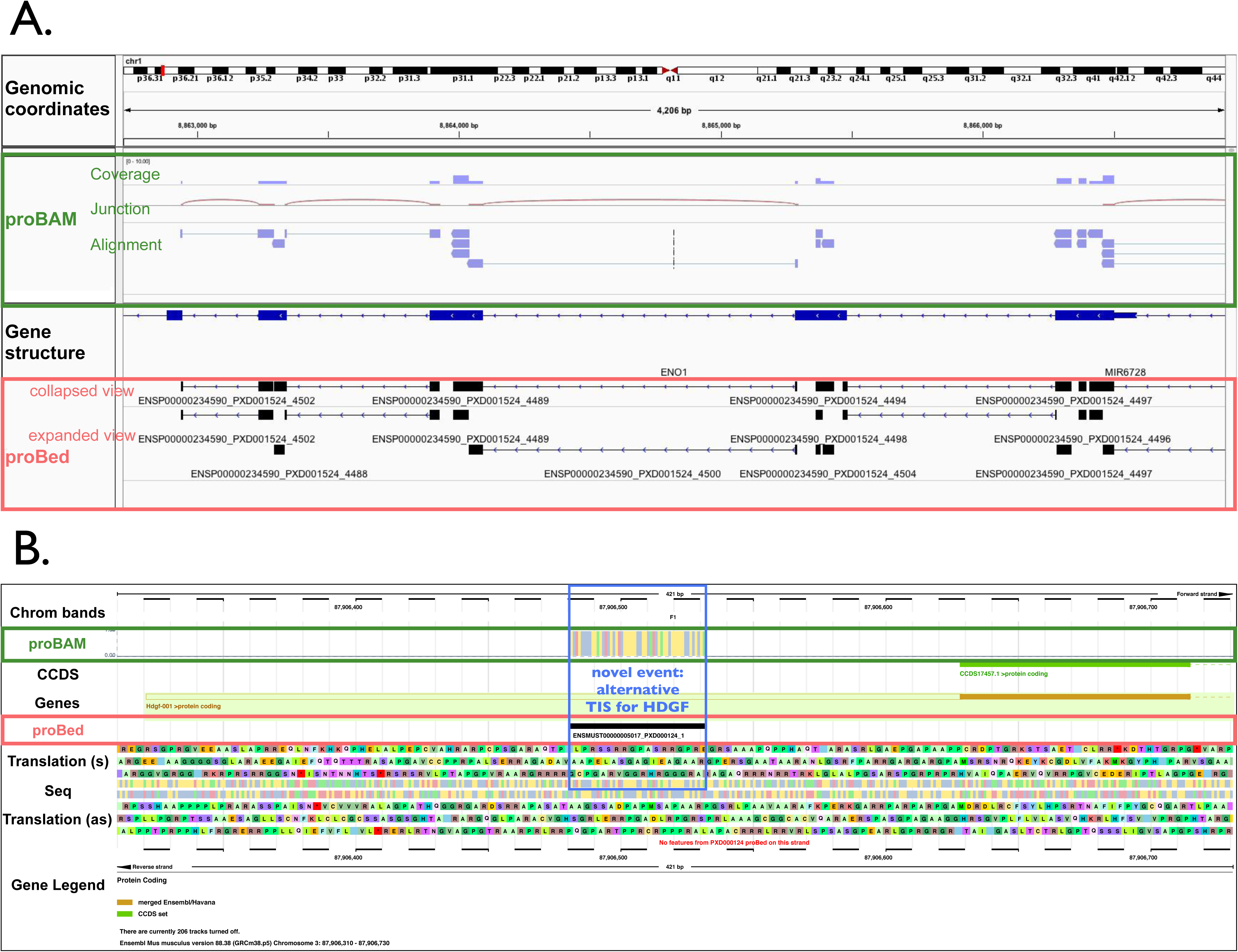
Visualization of proBAM and proBed files in genome browsers. a) IGV visualisation: proBAM (green box) and proBed (red box) files coming from the same dataset (accession number PXD001524 in the PRIDE database). proBed files are usually loaded as annotation tracks in IGV whereas proBAM files are loaded in the mapping section. b) Ensembl visualization: proBAM (green box) and proBed (red box) files derived from the same dataset (accession number PXD000124) illustrating a novel translational event. The N-terminal proteomics identification result points to an alternative translation initiation site (TIS) for the gene HDGF at a near-cognate start-site located in the 5’-UTR of the transcript (blue box).

The proBed format, on the other hand, is more tailored for storing only the final results of a given proteogenomics analysis, without providing the full details. The BED format is commonly used to represent genomic features. Thus, proBed stores browser track information at the PSM and/or peptide level mainly for visualisation purposes. As a key point, proBed files can be converted to BigBed [32], a binary format based on BED, which represents a feasible way to store the same information present in BED as compressed binary files, and is the final routinely used format as annotation tracks. It should be noted that a proBAM to proBed conversion should be possible, and vice versa. However, “null” values for some of the Tags would be logically expected for the mapping from proBed to proBAM.

### Software implementations

Both proBAM and proBed are fully compatible out-of-the-box with existing tools designed for the original SAM/BAM and BED files. Therefore, existing popular tools in the genomics community can readily be applied to read, merge and visualize these formats (Table 1). As mentioned already, several stand-alone and web genome browsers are available to visualize these formats e.g. UCSC browser, Ensembl, Integrative Genomics Viewer, and JBrowse.

**Table1.**
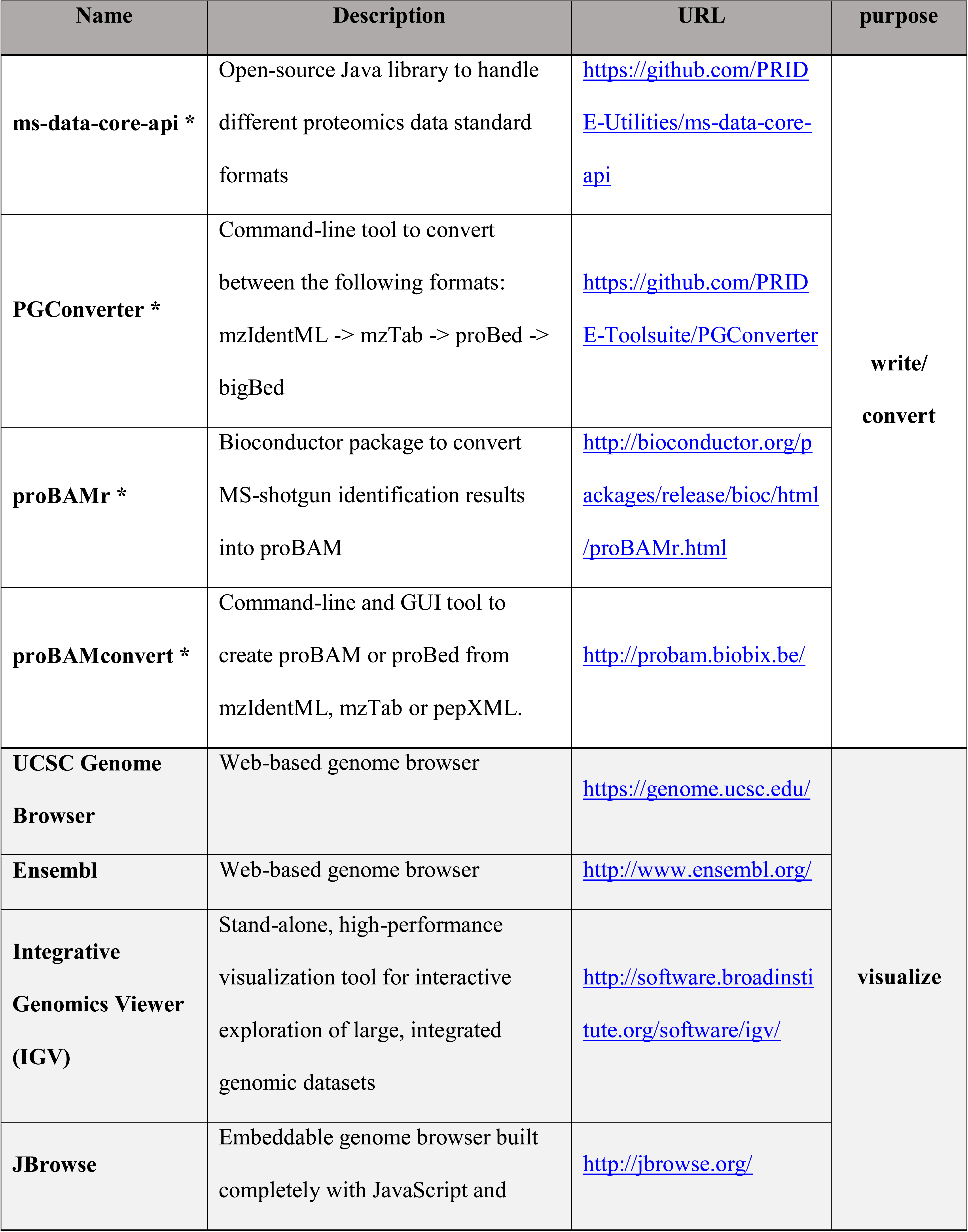
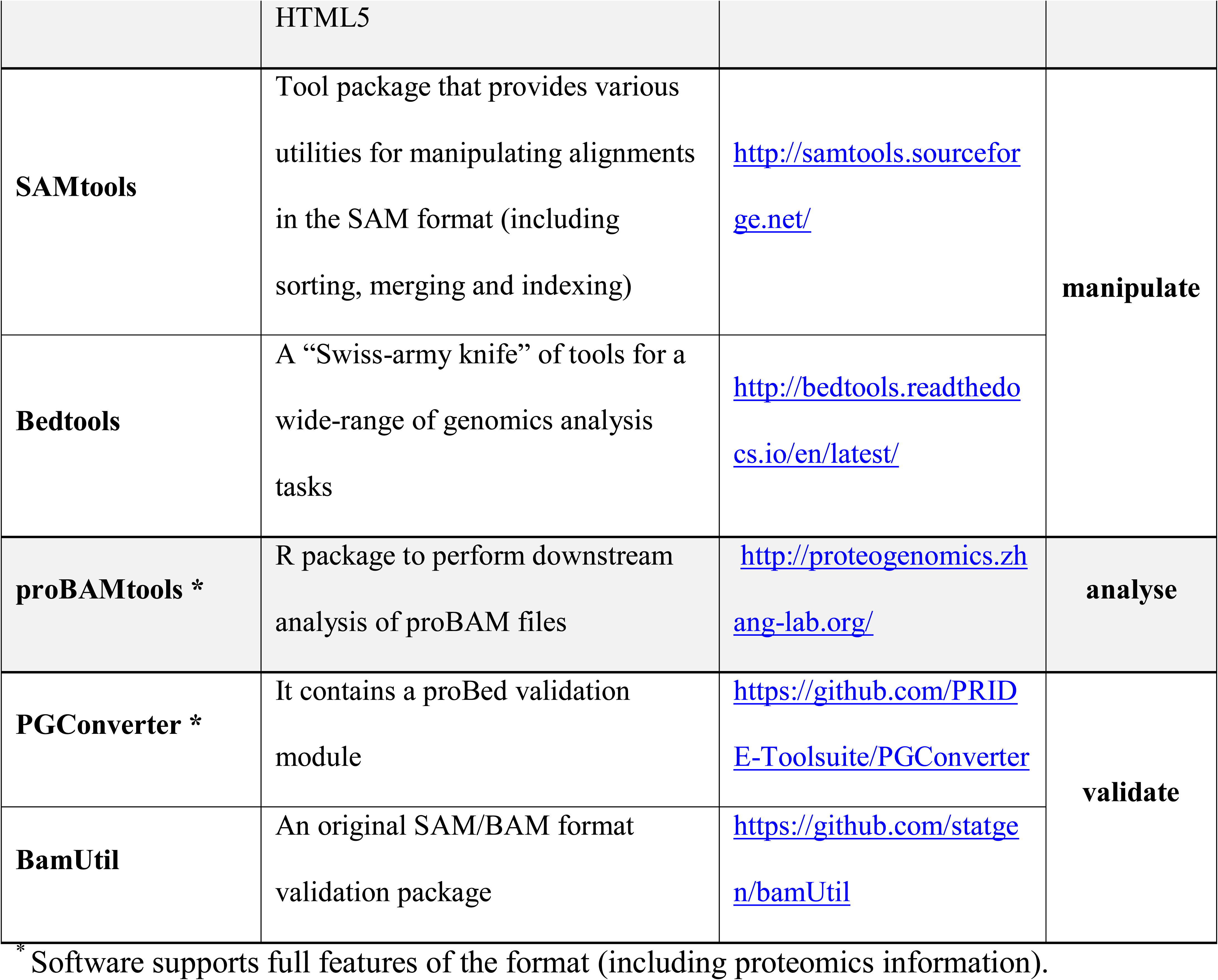
Existing software implementations of the proBAM and proBed formats (by June 2017).

Routinely used command line tools as SAMtools allow to manipulate (index, merge, sort) alignments in proBAM. Bedtools, seen as the “Swiss-army knife” tools for a wide-range of genomic analysis tasks, allows similar actions to both formats, including among others, intersection, merging, count, shuffling and conversion functionality. With the UCSC ‘bedToBigBed’ converter tool (http://hgdownload.soe.ucsc.edu/admin/exe/), one can also convert the proBed to bigBed. In this context, it is important to note that bedToBigBed version 2.87 is highlighted in the proBed format specification as the reliable version that can be used to create bigBed files coming from proBed (version 1.0) files.

There is also software specifically written for proBAM and proBed, supporting all the proteomics related features. In fact, proteogenomics data encoded in the PSI standard formats mzIdentML and mzTab can be converted into proBAM and proBed, although it should be noted that the representation for proteogenomics data in mzIdentML has only been formalized recently [33]. In this context, first of all, the open-source Java library ms-data-core-api, created to handle different proteomics file formats using the same interface, can be used to write proBed [34]. A Java command line tool, PGConverter (https://github.com/PRIDE-Toolsuite/PGConverter), is also able to convert from mzIdentML and mzTab to proBed and bigBed. Analogously, several tools are available to write proBAM files, such as the Bioconductor proBAMr package. An additional R package, called proBAMtools, is also available to analyze fully exported MS-based proteomics results in proBAM [31]. proBAMtools was specifically designed to perform various analyses using proBAM files, including functions for genome-based proteomics data interpretation, protein and gene inference, count-based quantification, and data integration. It also provides a function to generate a peptide-based proBAM file coming from a PSM-based one.

ProBAMconvert is another intuitive tool that enables the conversion from mzIdentML, mzTab and pepXML (another popular proteomics open format) [35] to both peptide- or PSM-based proBAM and proBed (http://probam.biobix.be) [36]. It is available as a command line interface (CLI) and a graphical user interface (GUI for Mac OS X, Windows and Linux). As CLI it is also wrapped in a Bioconda package (https://bioconda.github.io/recipes/probamconvert/README.html) and in a Galaxy tool, available from the public test toolshed (https://testtoolshed.g2.bx.psu.edu/view/galaxyp/probamconvert). The PGConverter tool also allows the validation of proBed files. For proBAM files, a validator is available that checks the validity of the original SAM/BAM format (https://github.com/statgen/bamUtil), although additional proteogenomics data verification still needs to be implemented.

## Discussion

We strongly believe that having available these two novel data formats (proBAM and proBed) constitutes an essential milestone for the continuous development of the field of proteogenomics. Successful promotion of proBAM and proBed requires support from software vendors, individual investigators, publishers, and data repositories. We will promote them following the typical channels used by the PSI. Therefore, further efforts will be focused on implementing these formats, not only using newly generated proteomics data but also on datasets already available in the public domain. In this context it is important to highlight that MS-based proteomics datasets are now routinely deposited in public repositories such as PRIDE [37], PeptideAtlas [38], MassIVE (https://massive.ucsd.edu) and jPOST [39] gathered in the ProteomeXchange Consortium (http://www.proteomechange.org [40]). In fact, an enormous amount of MS data is available in the public domain that can be used for proteogenomics studies, something that it is increasingly happening [41, 42]. The PRIDE database, located in the European Bioinformatics Institute (EMBL-EBI), plans to fully implement proBed in the coming months, facilitating the integration and visualisation of public proteomics data in Ensembl. In this context, it is also important to note that proBAM files generated from several large proteomics datasets have been already preloaded in a JBrowse-based genome browser (http://proteogenomics.zhang-lab.org/), facilitating the access to this data to a broader audience, both within and outside the proteomics community.

Additionally, we have already been actively pushing the use of these formats in big Consortia such as Clinical Proteomic Tumor Analysis Consortium (CPTAC). We hope the data released by such projects will inspire new tools that support these two formats. We expect that their existence will facilitate integration, visualization and exchange throughout both the proteomics and genomics communities, and will help multiple proteogenomics endeavours in trying to interpret proteomics results and/or refine gene model annotation by means of protein level validation.

The formats will be fully maintained by the PSI group using the strategy applied for all existing standard formats. If changes in the formats were needed that would not make them compatible with existing software, the formats would change their version number, and they would re-enter a new round of review in the PSI document process. Some future possible expansions for both formats could consider extended mechanisms to encode quantitative proteomics data. There is a mechanism to report PSM counts in proBed, but it is limited at present. Additionally, PSM counts can be calculated, at both gene and protein levels, from proBAM files. In the future, quantification support could be extended to additional workflows (e.g. intensity-based approaches).

We also highly encourage proteogenomics data providers to report PSMs to these two formats as part of their data exports, so it can be visualized by genome browsers directly and it is possible to re-analyse it within a genome context. We expect that the release and usage of proBed and proBAM will increase data sharing and integration between both the genomics and proteomics communities. The PSI remains a free and open consortium of interested parties, and we encourage critical feedback, suggestions and contributions *via* attendance at a PSI annual meeting, conference calls or our mailing lists (see http://www.psidev.info/).

## Acknowledgements

J.A.V., T.T., A.R.J. and F.G. want to acknowledge funding by the BBSRC grants “ProteoGenomics” [grant number BB/L024225/1] and “PROCESS” [grant number BB/K01997X/1], and A.R.J. wants to acknowledge BBSRC grant BB/L005239/1. G.M. is a Fellow of the Research Foundation – Flanders (FWO-Vlaanderen) [G.M.,12A7813N]. X.W. and B.Z. are supported by National Cancer Institute award U24CA159988 and U24CA210954. E.W.D. acknowledges funding from NIGMS grant number R01GM087221 and NIBIB grant number U54EB020406. D.F. is supported by National Cancer Institute award U24CA210972 and by contract 13XS068 from Leidos Biomedical Research, Inc. Finally, the colleagues in the Proteomics Standards Initiative, including the reviewers of the proBAM and proBed format specifications in the PSI document process, are acknowledged for helpful discussions and feedback. We also want to thank Andy Yates (Ensembl team), for his useful comments.

The author(s) declare(s) that they have no competing interests.

